# The Role of Context Conditioning in the Reinstatement of Responding to an Alcohol-Predictive Conditioned Stimulus

**DOI:** 10.1101/2021.07.19.450216

**Authors:** Mandy Rita LeCocq, Sophie Sun, Nadia Chaudhri

## Abstract

Re-exposure to an unconditioned stimulus (US) can reinstate extinguished conditioned responding elicited by a conditioned stimulus (CS). We tested the hypothesis that the reinstatement of responding to an appetitive CS is driven by an excitatory association formed between the US and the context that the US was ingested in during US re-exposure. Male, Long-Evans rats were acclimated to drinking alcohol (15%, *v*/*v*) in the home-cage, then trained to associate an auditory CS with an alcohol-US that was delivered into a fluid port for oral intake. During subsequent extinction sessions, the CS was presented as before, but without alcohol. After extinction, rats were re-exposed to alcohol as in training, but without the CS (alcohol re-exposure). 24 h later at test, the CS was presented as in training, but without alcohol. First, we tested the effect of extinguishing the context-alcohol association, formed during alcohol re-exposure, on reinstatement. Conducting four context extinction sessions across four days (spaced extinction) after the alcohol re-exposure session did not impact reinstatement. However, four context extinction sessions conducted across two days (massed extinction) prevented reinstatement. Next, we conducted alcohol re-exposure in a context that either differed from, or was the same as, the test context. One alcohol re-exposure session in a different context did not affect reinstatement, however, three alcohol re-exposure sessions in a different context significantly reduced reinstatement during the first CS trial. These results partially support the view that a context-US association formed during US re-exposure drives the reinstatement of responding to an appetitive, alcohol-predictive CS.

## Introduction

An important aspect of animal behaviour is the ability to associate a neutral, environmental stimulus with a salient, unconditioned stimulus (US). As a result of learning this predictive relationship, the environmental stimulus becomes a conditioned stimulus (CS) that can elicit conditioned responses. Conditioned responses allow animals to respond advantageously to environmental cues that predict appetitive stimuli (e.g., food, water) and aversive stimuli (e.g., predators, malaise). Importantly, in scenarios where the expected US stops occurring, animals learn to inhibit conditioned responding. This inhibition of responding, however, is not permanent and certain conditions can prompt a return of responding to the extinguished CS. For example, re-exposure to the US after extinction can reinstate conditioned responding to the CS (Bouton & Bolles, 1979; Rescorla & Heth, 1975; Stewart, 1984). Reinstatement is a fundamental phenomenon that is used to study learning and memory processes, including those related to extinction (Bouton & King, 1983; Monfils, Cowansage, Klann, & Ledoux, 2009). Reinstatement also has practical applications: it provides insight into how maladaptive reactions to environmental cues contribute to drug use and relapse (Jaffe, Cascella, Kumor, & Sherer, 1989; Rubonis et al., 1994) or post-traumatic stress disorder (Le Dorze & Gisquet-Verrier, 2016; Noble et al., 2017; Zuj, Palmer, Malhi, Bryant, & Felmingham, 2018) in clinical populations. As such, it is critical to understand the psychological processes that underlie the reinstatement effect.

Reinstatement has been predominantly studied using an aversive Pavlovian conditioning procedure in which rats are trained to associate an auditory CS with a foot shock-US, then the shock-US is withheld to extinguish conditioned responding. Next, rats receive unsignalled re-exposure to the shock-US, which reinstates conditioned responding to the CS at test 24 h later. This reinstatement of conditioned responding to an aversive CS, however, is only observed under specific conditions. For example, US re-exposure must be conducted in the same context that the subsequent reinstatement test occurs in. When re-exposure to the shock-US occurs in a context that differs from the test context, then reinstatement of the CS-evoked response does not occur (Bouton, 1984; Bouton & Bolles, 1979; Bouton & King, 1983, 1986; Frohardt, Guarraci, & Bouton, 2000; Westbrook, Iordanova, McNally, Richardson, & Harris, 2002; Wilson, Brooks, & Bouton, 1995). Additionally, if shock re-exposure is followed by sessions of repeated exposure to the context in which US re-exposure was conducted, then reinstatement is not observed (Bouton & Bolles, 1979). Reinstatement of conditioned responding to appetitive stimuli also requires these specific conditions. Re-exposure to a food-pellet in a context that differs from the test context reduces reinstatement of a CS-evoked response (Bouton & Peck, 1989) and of an operant lever pressing response (Baker, Steinwald, & Bouton, 1991). Furthermore, repeated exposure to the context that re-exposure to a food-pellet occurred in also attenuates reinstatement of an operant lever-pressing response (Baker et al., 1991). Based on these findings, it has been suggested that US re-exposure produces an excitatory association between the context and US. This ‘context-US association’ is proposed to summate with the residual, predictive value of the CS that survives extinction to drive reinstatement (Bouton & Peck, 1989; Bouton, Rosengard, Achenbach, Peck, & Brooks, 1993; Westbrook et al., 2002). This hypothesis is further supported by context preference tests conducted after US re-exposure. Rats avoid contexts associated with the shock-US delivered during US re-exposure, indicating that the context becomes associated with the shock-US (Bouton, 1984; Bouton & King, 1983).

The reinstatement of conditioned responding observed 24 h after US re-exposure is a reliable phenomenon observed in a variety of aversive and appetitive conditioning procedures. Re-exposure to an aversive US can reinstate various conditioned responses, such as freezing (Richardson, Duffield, Bailey, & Westbrook, 1999; Westbrook et al., 2002), suppression of an operant response (Bouton & Bolles, 1979; Bouton & King, 1983; Frohardt et al., 2000; Rescorla & Cunningham, 1977; Rescorla & Heth, 1975; Waddell, Morris, & Bouton, 2006; Wilson et al., 1995), fear-potentiated startle (Gewirtz, Falls, & Davis, 1997; Waddell, Bouton, & Falls, 2008), and taste aversion (Schachtman, Brown, & Miller, 1985). In appetitive conditioning, re-exposure to a food-pellet-US reinstated conditioned responding to an extinguished CS 24 h later (Bouton & Peck, 1989; Bouton et al., 1993). In an operant task, re-exposure to a food-pellet reinstated an extinguished operant lever-pressing response 24 h later at test (Baker et al., 1991). Conditioned responding for psychoactive substances can also be reinstated. After an established conditioned place preference for cocaine or methamphetamine was extinguished, re-exposure to the respective drug reinstated a preference for the drug-paired chamber 48 h later, relative to a chamber in which the drug was never delivered (Abulseoud, Miller, Wu, Choi, & Holschneider, 2012; Barbosa-Méndez, Matus-Ortega, Jacinto-Gutiérrez, & Salazar-Juárez, 2018; Hammad, Alasmari, Althobaiti, & Sari, 2017). Therefore, reinstatement is a general phenomenon that occurs in a wide range of learning paradigms.

We recently extended this literature by demonstrating the reinstatement of responding to a CS that predicted an appetitive, psychoactive substance, alcohol (LeCocq, Lahlou, Chahine, Padillo, & Chaudhri, 2018). In our task, following the acquisition and extinction of responding to an alcohol-predictive CS, re-exposure to an alcohol-US significantly reinstated responding to the CS 24 h later. Interestingly, a control group that received water instead of alcohol during US re-exposure showed similar reinstatement to that produced by alcohol. In a separate experiment where the control fluid was lemon-flavored water, making it distinct from alcohol, reinstatement was greater following re-exposure to the alcohol-US compared to the control fluid. In contrast, reinstatement was not observed when US re-exposure was delivered via systemic alcohol injection so that the pharmacological effects of alcohol were experienced without ingesting alcohol. These results provide new evidence of reinstatement to a CS that predicts a psychoactive substance, which occurs 24 h after exposure to the psychoactive substance. The present research was aimed at understanding the psychological processes underpinning this reinstatement effect.

As described above, research using aversive Pavlovian conditioning procedures suggests that a context-US association formed during US re-exposure summates with the residual, predictive value of the CS that survives extinction to drive reinstatement (Bouton & Peck, 1989; Bouton et al., 1993; Westbrook et al., 2002). However, there are fundamental differences in how associations are formed with aversive and appetitive stimuli that may result in different processes underlying reinstatement to an alcohol-predictive CS. First, the rate at which conditioned responses are acquired differs. Although most studies conduct more than one conditioning trial, subjects can associate a CS and an aversive US after just one pairing (Fanselow, 1990; Maes & Vossen, 1992; Mahoney & Ayres, 1976; Swank & Bernstein, 1994). Conversely, appetitive conditioning tasks typically require a greater number of CS-US pairings to form an association and evoke a conditioned response (Austen & Sanderson, 2020; Harris, 2011). Aversive and appetitive tasks also differ in how subjects interact with the US. In many aversive tasks, the US (e.g., foot shock) is experimentally delivered and results in a passive, non-voluntary experience (Bouton & Bolles, 1979; Schachtman et al., 1985). In appetitive tasks, the US is experimentally delivered (e.g., drops of alcohol, food pellets), but subjects must approach and ingest the US, which are voluntary behaviours (Bouton et al., 1993; Valyear et al., 2020; Villaruel et al., 2018). Moreover, appetitive tasks often require subjects to engage in a consummatory response to ingest the US, unless the US is delivered via systemic injection (Parker & Mcdonald, 2000), intragastric intubation (Sclafani, Fanizza, & Azzara, 1999) or intrajugular catheter (Shaham & Stewart, 1994; Uslaner, Acerbo, Jones, & Robinson, 2006). In the latter cases, the route of US administration is non-voluntary, making these procedures more akin to passive aversive conditioning. Given these fundamental differences in appetitive and aversive conditioning, the extent to which a context-US association drives reinstatement to an alcohol-predictive CS is unknown.

We investigated the psychological processes underlying the reinstatement of responding to an alcohol-predictive CS using two distinct behavioural manipulations that assessed the role of a context-US association in reinstatement. In Experiment 1, we tested the effect of extinguishing the context-alcohol association formed during alcohol re-exposure on reinstatement. In Experiment 2, we tested the effect of conducting alcohol re-exposure in a context that differed from the subsequent test context on reinstatement.

## Methodology

### Subjects

Male, Long-Evans rats (Envigo, Indianapolis, IN; 220 – 240 g on arrival) were pair-housed upon arrival and left unhandled for three days. Next, rats were single housed and handled for five days before experimental procedures began. Cages contained beta chip bedding (Aspen Sani chips, Envigo), a nylabone toy (Nylabones, Bio-Serv), a polycarbonate tunnel (Rat retreats, Bio-Serv), and shredded paper. Unrestricted access to water and rat chow (Purina Agribrands, Charles River) was provided throughout the experiment. Cages were held in a temperature-(21° C) and humidity-controlled (40 - 50%) colony room that was on a 12 h light/dark cycle (lights on at 0700 h; all experimental procedures occurred during the light phase). All procedures followed the guidelines of the Canadian Council on Animal Care and were approved by the Concordia University Animal Research Ethics Committee.

### Materials

#### Apparatus

Behavioural procedures were conducted in conditioning chambers (ENV-009A; Med Associates Inc., St-Albans, VT) that were enclosed in sound-attenuating, ventilated melamine cubicles (made in house). Chambers included a Plexiglass front door, back-wall and ceiling, stainless steel sidewalls, and a metal bar floor (ENV-009A-GF). A house light (ENV-215M; 75W, 100 mA) and a white-noise generator (ENV-225SM, 5 dB above background noise) were mounted on the upper left chamber wall. A dual-cup, fluid port (ENV-200R3AM) was located off-centred, on the lower right chamber wall. Alcohol was delivered to the fluid port via polyethylene tubing from a 20 mL syringe mounted on a syringe pump (PHM-100, 3.33 RPM) located outside the melamine cubicles. Port entries were measured via interruptions of an infrared beam across the entrance of the fluid port. Med PC IV software on a PC controlled stimulus delivery and recorded behavioural responses.

#### Solutions

Alcohol solutions (5%, 10%, 15% *v*/*v*) were prepared by diluting 95% ethanol in tap water. Odours for context configurations were prepared by diluting 10% *v*/*v* lemon oil (lemon odour; Sigma Aldrich, CAS# 8008-56-8) or 10% *v*/*v* benzaldehyde (almond odour; bought inhouse at Concordia University chemistry store) in tap water.

### Behavioural procedures

#### Intermittent alcohol access

Rats received 15 sessions of intermittent access to 15% (*v*/*v*) alcohol in the home-cage to induce high levels of alcohol intake (Carnicella, Ron, & Barak, 2014; Sparks, Sciascia, Ayorech, & Chaudhri, 2013; Wise, 1973). During alcohol access sessions (Monday, Wednesday, Friday), rats were first weighed, then pre-weighed 100 mL graduated cylinders containing alcohol and pre-weighed water bottles were inserted into home-cages via the cage lid. 24 h later, the alcohol cylinders were replaced with water cylinders (Tuesday, Thursday, Saturday, Sunday). Cylinder placement was randomized across the left and right sides of the cage lid to mitigate the impact of side preference on drinking. Cylinders and water bottles were weighed after each 24 h session to calculate fluid intake. Spillage was accounted for by subtracting the average amount of fluid lost from bottles placed on two empty control cages from the corresponding session data.

The unsweetened alcohol solution used in this procedure produces variable levels of alcohol intake (Cofresi et al., 2017; LeCocq et al., 2018; Sparks et al., 2013). For rats with low alcohol intake, we reduced the alcohol concentration to encourage drinking (Cofresi et al., 2019, 2017). Rats that drank < 1 g/kg [g of alcohol consumed/kg of body weight] on three consecutive sessions subsequently received 5% (*v*/*v*) alcohol on session 5 (Experiment 1, *n* = 7) or on session 4 (Experiment 2, *n* = 2). Once 1 g/kg was obtained for two consecutive sessions, the alcohol concentration was increased to 10% (*v*/*v*). When 1 g/kg was obtained for two consecutive sessions the alcohol concentration was increased to 15% (*v*/*v*). Rats (*n* = 1) that drank < 1 g/kg for three consecutive sessions after session 6 subsequently received 10% (*v*/*v*) alcohol until 1 g/kg was obtained for two consecutive sessions and then the alcohol concentration was increased to 15% (*v*/*v*). The concentration of alcohol that rats received on the last intermittent alcohol access session was the alcohol concentration that they received during Pavlovian conditioning and alcohol re-exposure. Supplementary Table 1 depicts the number of rats that received 5% and 10% alcohol during subsequent training sessions.

#### Habituation

Two habituation sessions were conducted during the last week of intermittent alcohol access on days with only access to water. In session 1, rats were brought to the experimental room, weighed, and left in their home cages for 20 min. In session 2, rats were weighed and then placed into a designated conditioning chamber for 20 min, during which the house lights were illuminated, and the total number of port entries made was counted.

#### Pavlovian conditioning

Pavlovian conditioning was conducted daily (Monday – Friday). Sessions began with a 2 min delay, after which the house lights were illuminated to signal the start of the session. In each session, eight trials of a 20 s continuous white-noise conditioned stimulus (CS) were paired with 10 s activation of the fluid pump to deliver 0.3 mL of alcohol into the fluid port (2.4 mL per session). Pump activation began 10 s after CS onset and co-terminated with the CS. Trials occurred on a variable time 240 s schedule (intertrial intervals: 120, 200, 210, 280, 310, 320 s; not including 20 s pre- and post-CS intervals). Total session length was 42 - 45 min. Fluid ports were checked at the end of each session to verify that the alcohol was ingested.

#### Extinction

Extinction sessions were conducted daily (Sunday – Saturday). Session parameters were identical to Pavlovian conditioning except that the CS was paired with activation of empty syringe pumps (i.e., alcohol was not delivered).

#### Alcohol re-exposure

During alcohol re-exposure, 2.4 mL of alcohol was delivered into the fluid port according to the same schedule as Pavlovian conditioning; however, the CS was not presented.

#### Reinstatement test

During the reinstatement test, the CS was presented as during Pavlovian conditioning, except that CS trials were paired with activation of empty syringe pumps (i.e., alcohol was not delivered). The experimental procedures for all experiments are illustrated in Table 1.

**Table 1.** Experimental Design

| Experiment 1A |  |  |  |  |  |
| --- | --- | --- | --- | --- | --- |
| Group | Conditioning<br>16 sessions | Extinction<sup>a</sup><br>7 sessions | Alcohol<br>re-exposure<br>1 session | Context exposure<br>4 sessions (1/day) | Test<br>1 session |
| No extinction | CS+US | CS | US | Alternate context | CS |
| No port | CS+US | CS | US | Conditioning chamber (no port) | CS |
| Context extinction | CS+US | CS | US | Conditioning chamber | CS |
| Experiment 1B |  |  |  |  |  |
| Group | Conditioning<br>2 sessions | Extinction<sup>a</sup><br>5 sessions | Alcohol<br>re-exposure<br>1 session | Context exposure<br>4 sessions (2/day) | Test<br>1 session |
| No extinction | CS+US | CS | US | Alternate context | CS |
| No US | CS+US | CS | No US | Conditioning chamber or alternate context | CS |
| Context extinction | CS+US | CS | US | Conditioning chamber | CS |
| Experiment 2A |  |  |  |  |  |
| Group | Conditioning<br>16 sessions | Extinction<sup>b</sup><br>7 sessions | Alcohol re-exposure<br>1 session | Test<br>1 session |  |
| Different | CS+US (Context A) | CS (Context A) | US (Context B) | CS (Context A) |  |
| Same | CS+US (Context A) | CS (Context A) | US (Context A) | CS (Context A) |  |
| Experiment 2B |  |  |  |  |  |
| Group | Conditioning<br>2 sessions | Extinction<sup>c</sup><br>5 sessions | Alcohol re-exposure<br>3 sessions | Test<br>1 session |  |
| Different | CS+US (Context A) | CS (Context A) | US (Context B) | CS (Context A) |  |
| Same | CS+US (Context A) | CS (Context A) | US (Context A) | CS (Context A) |  |

<sup>a</sup> Experiment 1A and 1B: Habituation to the alternate context was conducted after every extinction session.
<sup>b</sup> Experiment 2A: Context B habituation sessions were conducted after extinction sessions 5 and 6.
<sup>c</sup> Experiment 2B: Context B habituation sessions were conducted after extinction sessions 3 and 4.

### Experiment 1A: The effect of spaced context extinction on reinstatement

We tested the effect of extinguishing the context-alcohol association formed during the alcohol re-exposure session on reinstatement. After intermittent alcohol access and habituation, rats (*n* = 36) received 16 Pavlovian conditioning sessions, then seven extinction sessions. Four hours after each extinction session, all rats were habituated to a covered plastic bucket containing Sani-chip bedding for 20 min (‘alternate context’). One alcohol re-exposure session was conducted 24 h after the last extinction session. Rats were then divided into three groups matched on body weight, ΔCS port entries, and total port entries across Pavlovian conditioning and extinction sessions.

Starting 24 h after alcohol re-exposure, each group received four context exposure sessions across four consecutive days, as this design has been shown to attenuate the reinstatement of operant responding for food-pellets (Baker et al., 1991). Rats in the ‘Context extinction’ group (*n* = 12) were placed into the conditioning chambers for daily context extinction sessions in which house light onset occurred after a 2 min delay, but no CS or alcohol were delivered. Rats in the ‘No port’ group (*n* = 12) received identical sessions; however, the fluid port was replaced with a metal panel. This control group was included to account for the possible effect on reinstatement of extinguishing consummatory port entry responses, rather than extinguishing the context-alcohol association. Rats in the ‘No extinction’ group (*n* = 12) were placed in the alternate context for the same duration as both other groups. At 24 h after the last context exposure session, all groups received a reinstatement test in the conditioning chambers.

### Experiment 1B: The effect of massed context extinction on reinstatement

Because all groups in Experiment 1A showed reinstated conditioned responding to the CS, we tested a massed context extinction design which has been shown to attenuate the reinstatement of conditioned responding to a shock-predictive CS (Bouton & Bolless, 1979). We reasoned that massed context extinction would be more effective than spaced context extinction at extinguishing the context-alcohol association, as massed extinction trials produce more robust extinction of conditioned responding to a CS than spaced trials (Cain, Blouin, & Barad, 2003). Rats from Experiment 1A (*n* = 35) received two Pavlovian conditioning sessions, then five extinction sessions with habituation to the alternate context occurring four hours after every extinction session. Rats were then divided into the same groups they were assigned in Experiment 1A. The ‘Context extinction’ (*n* = 12) and ‘No extinction’ (*n* = 11) groups received one alcohol re-exposure session followed by context extinction or exposure to the alsternate context, respectively, as in Experiment 1A. Rats that were previously in the ‘No port’ group were included in a ‘No US’ control group (*n* = 12) to assess if spontaneous recovery might contribute to responding at test. Rats in this group were placed in conditioning chambers with the house light illuminated but did not get alcohol during the alcohol re-exposure session. Next, half the rats received exposure to the conditioning chambers while the remainder received exposure to the alternate context. In this experiment, context exposure sessions were conducted across two days for all groups. The first session occurred at the time that previous phases of training had been conducted (1300 h) and the second session occurred at 1700 h. All groups received a test for reinstatement in the conditioning chambers 24 h after the last context exposure session.

### Experiment 2A: The effect of one alcohol re-exposure session in a different context on reinstatement

We examined the effect of conducting alcohol re-exposure in a context that differed from the reinstatement test context on reinstatement. After intermittent alcohol access and habituation, rats (*n* = 28) received 16 Pavlovian conditioning sessions, then seven extinction sessions that were all conducted in Context A. Four hours after the second-to-last and the third-to-last extinction sessions, rats were habituated to Context B for 20 min. Context A and B configurations were counterbalanced across Context Type 1 (grid floor, almond scent, transparent doors) and Context Type 2 (Perspex floor, lemon scent, black opaque doors). After extinction, rats were divided into two groups that were matched on body weight, ΔCS port entries, and total port entries, made across Pavlovian conditioning and extinction sessions. The ‘Same’ group (*n* = 14) received one alcohol re-exposure session in Context A, while the ‘Different’ group (*n* = 14) received one alcohol re-exposure session in Context B. At 24 h later, both groups were tested for reinstatement in Context A.

### Experiment 2B: The effect of three alcohol re-exposure sessions in a different context on reinstatement

Because in Experiment 2A both groups showed reinstatement, we tested the possibility that one alcohol re-exposure session was not sufficient for rats to associate the different context with the alcohol-US. Consequently, we examined the effect of three US re-exposure sessions in a different context on reinstatement. Rats from Experiment 2A (*n* = 28) received two Pavlovian conditioning sessions, then five extinction sessions in Context A. The third-to-last and second-to-last extinction sessions were followed by habituation to Context B, as described above. Rats were divided into the ‘Different’ (*n* = 14) and ‘Same’ (*n* = 14) groups then received three alcohol re-exposure sessions, as per their Experiment 2A group assignments. At 24 h later, both groups were tested for reinstatement in Context A.

### Data Management

#### Exclusion criteria

Rats did not transition from intermittent alcohol access to behavioural training if they drank <1 g/kg averaged across the last three sessions of intermittent alcohol access (Experiment 1; *n* = 1: Experiment 2; *n* = 0), received 10% alcohol on session 13 and onwards (Experiment 1; *n* = 1: Experiment 2; *n* = 0), or displayed aggressive behaviour (Experiment 1; *n* = 1: Experiment 2; *n* = *0*).

Following training, we used a behavioural criterion based on the probability of making a CS port entry [# of trials with ≥1 CS port entry / # of CS trials) * 100] to evaluate if rats had learned to associate the CS with alcohol. Rats with a probability score of ≤ 70% averaged across the last two Pavlovian conditioning sessions were removed from statistical analyses (Experiment 1A, *n* = 3; Experiment 1B, *n* = 4; Experiment 2A, *n* = 3; Experiment 2B, *n* = 3). Rats with a probability score of ≥ 60% averaged across the last two extinction sessions were also removed from statistical analyses (Experiment 1A, *n* = 2; Experiment 1B, *n* = 3; Experiment 2A, *n* = 3; Experiment 2B, *n* = 2). One rat from Experiment 1A became highly aggressive during training and was removed from the study. Table 2 depicts initial and final sample sizes for all experiments.

**Table 2.** Sample Size Across Experimental Phases

| Exp. | Intermittent alcohol access |  | Conditioning |  | Extinction |  | Final sample size |
| --- | --- | --- | --- | --- | --- | --- | --- |
|  | Initial | Dropped | Initial | Dropped | Initial | Dropped |  |
| 1A | $n = 39$ | $n = 3^a$ | Context extinction $n = 12$<br>No extinction $n = 12$<br>No port $n = 12$ | $n = 3$ | Context extinction $n = 11$<br>No extinction $n = 11^b$<br>No port $n = 11$ | $n = 3$ | Context extinction $n = 10$<br>No extinction $n = 9$<br>No port $n = 11$ |
| 1B | N/A | N/A | Context extinction $n = 12$<br>No extinction $n = 11$<br>No US $n = 12$ | $n = 4$ | Context extinction $n = 11$<br>No extinction $n = 9$<br>No US $n = 11$ | $n = 3$ | Context extinction $n = 10$<br>No extinction $n = 9$<br>No US $n = 9$ |
| 2A | $n = 28$ | $n = 0$ | Same $n = 14$<br>Different $n = 14$ | $n = 3$ | Same $n = 12$<br>Different $n = 13$ | $n = 3$ | Same $n = 10$<br>Different $n = 12$ |
| 2B | N/A | N/A | Same $n = 14$<br>Different $n = 14$ | $n = 3$ | Same $n = 12$<br>Different $n = 13$ | $n = 2$ | Same $n = 12$<br>Different $n = 11$ |

<sup>a</sup> Rats dropped because of $<1$ g/kg across last three intermittent access sessions ( $n = 1$ ), received 10% alcohol on session 13 or onwards ( $n = 1$ ), aggressive ( $n = 1$ ).
<sup>b</sup> Rat dropped because aggressive ( $n = 1$ ).

#### Variables

Our dependent variables were ΔCS port entries (CS port entries minus port entries during the 20 s pre-CS interval), total duration of CS port entries (length of time (s) spent in the fluid port summed across CS trials), and average latency to make a CS port entry (time (s) to initiate the first port entry during each CS trial averaged across CS trials). If a port entry was not made during a CS trial, a latency value of 20 s was used (LeCocq et al., 2018; Millan, Reese, Grossman, Chaudhri, & Janak, 2015).

### Statistical Analyses

Responding at test was compared to an extinction baseline obtained by averaging data across the last two extinction sessions. Data from the context extinction sessions in Experiment 1A and 1B were analyzed using a repeated measures analysis of variance (ANOVA). Data from the reinstatement test were analyzed using a mixed Phase x Group ANOVA and a mixed Trial x Group ANOVA. The Huynh-Feldt correction was applied when Mauchly’s test of sphericity was violated. All post-hoc analyses were corrected for multiple comparisons with the Bonferroni adjustment. Statistical analyses were conducted with IBM SPSS (Version 23; IBM Corp., Armonk, NY) and evaluated using a statistical significance level of *p* < 0.05. Graphs were created with Graphpad Prism (Version 7; La Jolla, CA).

## Results

Alcohol intake increased across intermittent alcohol access sessions in Experiment 1 and Experiment 2 (Supplementary Figure 1). Rats learned to associate the CS with alcohol, as ΔCS port entries significantly increased across Pavlovian conditioning sessions in Experiments 1 and 2 (Supplementary Figure 2). ΔCS port entries significantly decreased across extinction sessions in Experiment 1 and 2 (Supplementary Figure 2).

### Experiment 1A: Spaced context extinction did not affect reinstatement

Following Pavlovian conditioning, extinction and alcohol re-exposure, rats received four daily sessions (i.e., spaced context extinction) of exposure to an alternate context (‘No extinction’), the conditioning chambers (‘Context extinction’), or the conditioning chambers without fluid ports (‘No port’). Total port entries made by the ‘Context extinction’ group showed a significant reduction (Figure 1A) across context extinction sessions [*F*_(3, 27)_ = 2.99, *p* = .048], suggesting that context extinction had occurred.

**Figure 1.**
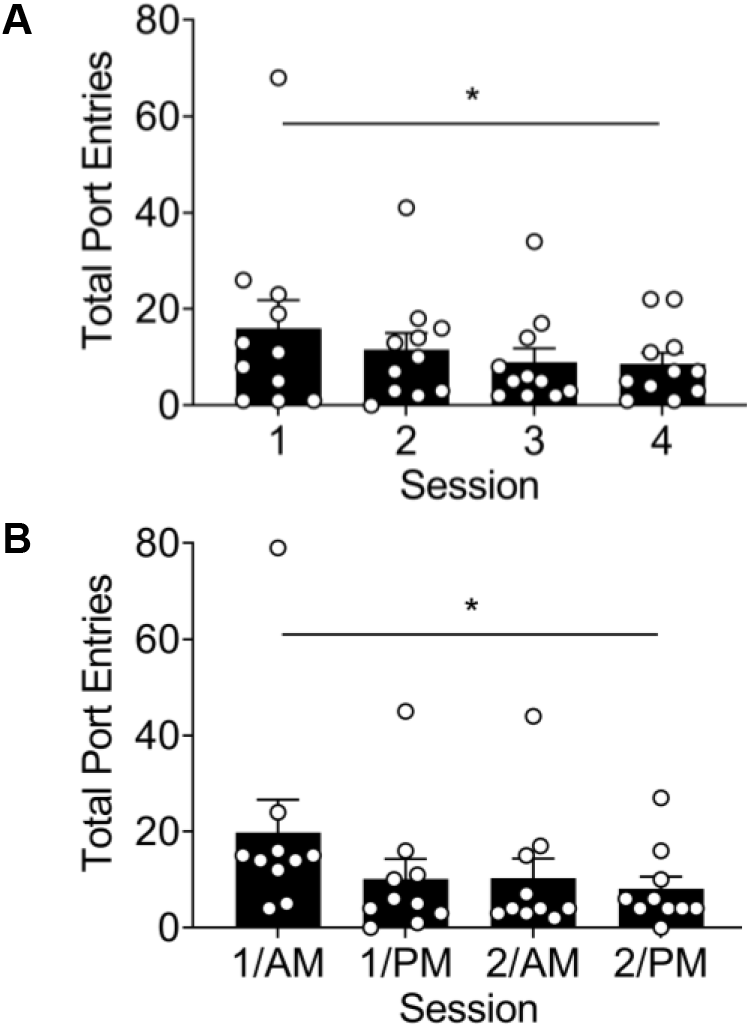
Conducting spaced or massed context extinction sessions significantly reduced total port entries across sessions. Data represent mean (± SEM) total port entries from rats in the ‘Context extinction’ group in **A** Experiment 1A (*n* = 10), and **B** Experiment 1B (*n* = 10). Open circles depict data of individual rats. * *p* < 0.05, main effect of Session (1 – 4)

Following spaced context exposure sessions, all groups reinstated to a similar degree. Relative to extinction, ΔCS port entries (Figure 2A) significantly increased at test [Phase: *F*_(1, 27)_ = 46.64, *p* < .001] in all three groups [Group: *F*_(2, 27)_ = 0.15, *p* = .865; Phase x Group: *F*_(2, 27)_ = 0.40, *p* = .673]. Similar effects were found in the total duration of CS port entries (Figure 2B) and the average latency to initiate the first CS port entry (Figure 2C). At test, all groups showed a significant increase in the duration of CS port entries [Phase: *F*_(1, 27)_ = 46.45, *p* < .001; Group: *F*_(2, 27)_ = 0.76, *p* = .480; Phase x Group: *F*_(2, 27)_ = 0.25, *p* = .785] and a significant decrease in latency to CS port entries [Phase: *F*_(1, 27)_ = 34.60, *p* < .001; Group: *F*_(2, 27)_ = 0.57, *p* = .574; Phase x Group: *F*_(2, 27)_ = 1.04, *p* = .367], relative to extinction.

**Figure 2.**
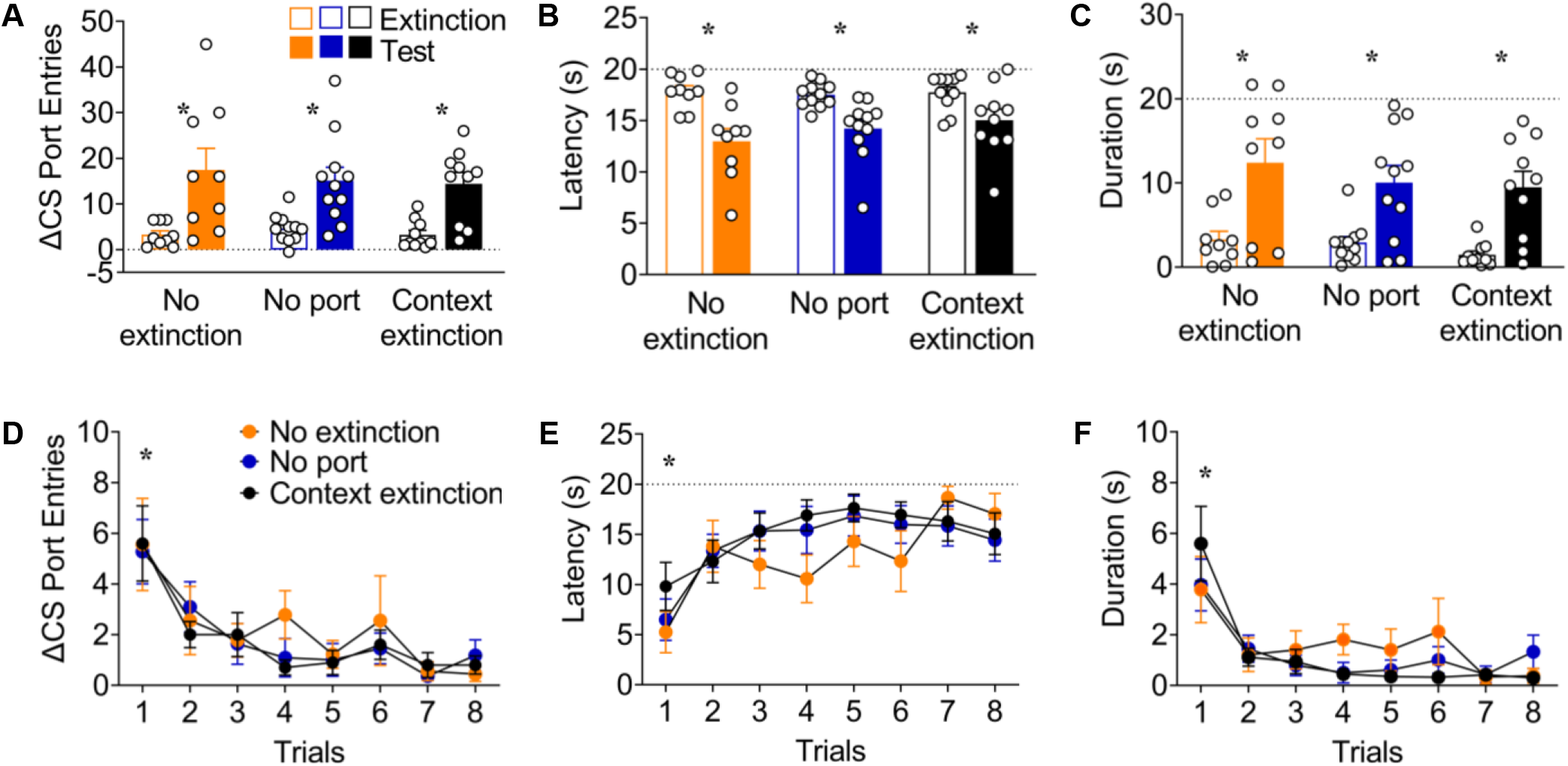
Conducting spaced context extinction after alcohol re-exposure did not reduce reinstatement. Data are from rats that received context exposure in an alternate context (No extinction; Orange; *n* = 9), the training context without fluid ports (No port; Blue; *n* = 11), or the training context (Context extinction; Black; *n* = 10). **A - C** Mean (± SEM) responding during extinction and test for **(a)** ΔCS port entries, **(b)** total duration of CS port entries, and **(c)** average latency to initiate the first CS port entry. **D - F** Mean (± SEM) responding across CS trials at test for **(d)** ΔCS port entries, **(e)** duration of CS port entries, and **(f)** latency to initiate the first CS port entry. Open circles depict data from individual rats. * *p* < 0.05, main effects of **A - C** Phase (Extinction vs Test) and **D - F** Trial (1 – 8)

To determine if spaced context extinction affected the pattern of responding at test, we analyzed port entries as a function of CS trial. ΔCS port entries (Figure 2D) significantly decreased across CS trials due to within-session extinction [Trial: *F*_(5.097, 137.626)_ = 10.82, *p* < .001] with no differences between groups [Group: *F*_(2, 27)_ = 0.20, *p* = .817; Trial x Group: *F*_(10.195, 137.626)_ = 0.38, *p* = .956]. The total duration of CS port entries (Figure 2E) significantly decreased across CS trials [Trial: *F*_(3.923, 105.909)_ = 12.14, *p* < .001] in all groups [Group: *F*_(2, 27)_ = 0.45, *p* = .641; Trial x Group: *F*_(7.845, 105.909)_ = 0.98, *p* = .454]. The average latency to initiate the first CS port entry (Figure 2F) significantly increased across CS trials [Trial: *F*_(7, 189)_ = 7.69, *p* < .001] in all groups [Group *F*_(2, 27)_ = 0.89, *p* = .421; Trial x Group: *F*_(14, 189)_ = 0.97, *p* = .476]. These results indicate that spaced context extinction had no effect on reinstatement, despite producing a significant reduction in total port entries across context extinction sessions.

### Experiment 1B: Massed context extinction prevented reinstatement

Following Pavlovian conditioning, extinction and alcohol re-exposure, rats from Experiment 1A received four sessions, conducted two times per day (i.e., massed context extinction), of exposure to an alternate context (‘No extinction’) or the conditioning chambers (‘Context extinction’). An additional control group (‘No US’) did not receive alcohol re-exposure and was then exposed to either the alternate context or the conditioning chambers. As responding in these two subgroups was similar at test, their data were collapsed into one group (i.e., ‘No US’) for statistical analyses. Total port entries made by the ‘Context extinction’ group showed a significant reduction (Figure 1B) across context extinction sessions [*F*_(1.1721, 15.488)_ = 5.15, *p* = .023] suggesting that context extinction had occurred.

Conducting massed context extinction sessions after alcohol re-exposure prevented reinstatement (Figure 3A). Relative to extinction, ΔCS port entries significantly increased at test [Phase: *F*_(1, 25)_ = 23.45, *p* < .001]. There was no overall effect group [Group: *F*_(2, 25)_ = 2.57, *p* = .097]; however, reinstatement differed between groups as a function of phase [Phase x Group: *F*_(2, 25)_ = 6.84, *p* = .004]. Post-hoc analyses revealed that the ‘No extinction’ group made significantly more ΔCS port entries at test relative to extinction (*p* < .001), whereas the ‘Context extinction’ (*p* = .068) and ‘No US’ (*p* = .466) groups did not. The total duration of CS port entries (Figure 3B) significantly increased at test [Phase: *F*_(1, 25)_ = 11.01, *p* = .003] in all three groups [Group: *F*_(2, 25)_ = 1.93, *p* = .166; Phase x Group: *F*_(2, 25)_ = 2.60, *p* = .094]. The latency to initiate the first CS port entry (Figure 3C) significantly decreased at test [Phase: *F*_(1, 25)_ = 20.94, *p* < .001]. There was no overall effect of group [Group: *F*_(2, 25)_ = 2.18, *p* = .134]; however, reinstatement differed between groups as a function of phase [Phase x Group: *F*_(2, 25)_ = 9.09, *p* = .001]. Post-hoc analyses revealed that the ‘No extinction’ group was significantly quicker to initiate a CS port entry at test relative to extinction (*p* < .001), whereas the ‘Context extinction’ (*p* = .740) and ‘No US’ (*p* = .138) groups were not.

**Figure 3.**
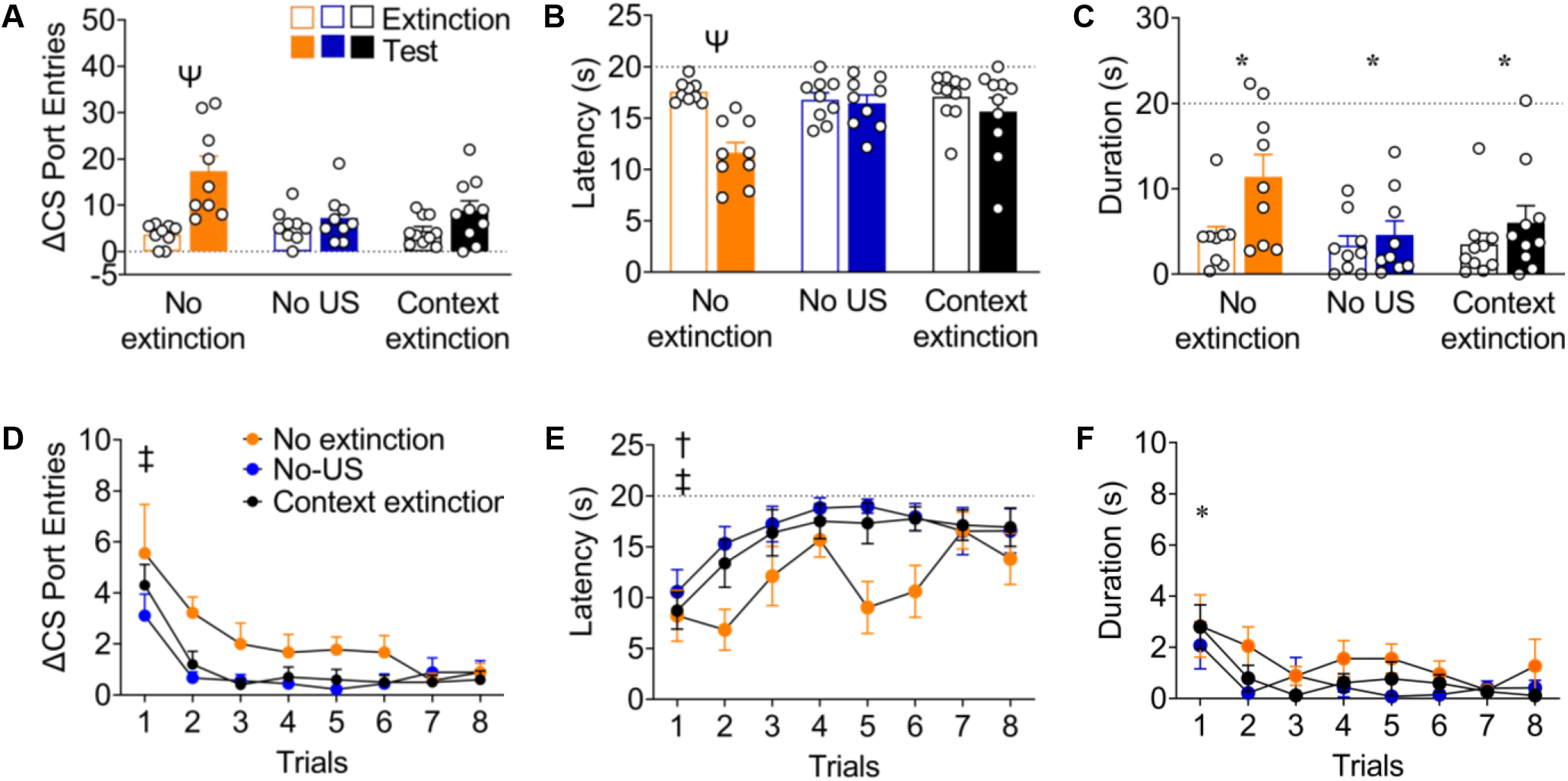
Conducting massed context extinction after alcohol re-exposure prevented reinstatement. Data are from rats that received context exposure in an alternate context (No extinction; Orange; *n* = 9), in the training context (Context extinction; Black; *n* = 10), or received no alcohol re-exposure then context exposure to the alternate or training context (No US; Blue; *n* = 9). **A – C** Mean (± SEM) responding during extinction and test for **(a)** ΔCS port entries, **(b)** total duration of CS port entries and **(c)** average latency to initiate the first CS port entry. **D - F** Mean (± SEM) responding across CS trials at test for **(d)** ΔCS port entries, **(e)** duration of CS port entries, and **(e)** latency to initiate the first CS port entry. Ψ *p* < 0.05, Phase x Group interaction post-hoc (Extinction vs. Test). * *p* < 0.05, main effects of **A – C** Phase (Extinction vs. Test) and **D – F** Trial (1 – 8). ‡ *p* < 0.05, main effect of Group post-hoc (No US vs. No extinction). † *p* < 0.05, main effect of Group post-hoc (Context extinction vs. No extinction).

In an analysis of responding as a function of CS trial at test, ΔCS port entries (Figure 3D) significantly decreased across CS trials [Trial: *F*_(2.875, 71.874)_ = 14.13, *p* < .001]. This effect differed as a function of group [Group: *F*_(2, 25)_ = 4.83, *p* = .017] with no significant interaction [Trial x Group: *F*_(5.750, 71.874)_ = 0.88, *p* = .512]. Post-hoc analyses revealed that the main effect of Group was driven by significantly more ΔCS port entries by the ‘No extinction’ group compared to the ‘No US’ group (*p* = .024). The ‘No extinction’ group made numerically more ΔCS port entries than the ‘Context extinction’ group; however, this difference did not reach statistical significance (*p* = .059). The ΔCS port entries by the ‘No US’ and ‘Context extinction’ groups did not differ (*p* = 1.0). The total duration of CS port entries (Figure 3E) significantly decreased across CS trials [Trial: *F*_(4.134, 103.358)_ = 5.07, *p* = .001] in all three groups [Group: *F*_(2, 25)_ = 2.83, *p* = .078; Trial x Group: *F*_(8.269, 103.358)_ = 0.54, *p* = .832]. The average latency to initiate the first CS port entry (Figure 3F) significantly increased across CS trials [Trial: *F*_(6.690, 167.240)_ = 1.98, *p* < .001]; however, this effect differed as a function of group [Group: *F*_(2, 25)_ = 5.40, *p* = .011] with no significant interaction [Trial x Group: *F*_(12.379, 167.240)_ = 1.13, *p* = .334]. Post-hoc analyses revealed that the main effect of Group was driven by the ‘No extinction’ group making CS port entries more quickly than the ‘Context extinction’ group (*p* = .047) and the ‘No US’ group (*p* = .016). There was no significant difference between the ‘Context extinction’ and the ‘No US’ groups (*p* = 1.0). These results, obtained across multiple measures of conditioning, show that conducting massed context extinction significantly attenuated reinstatement.

### Experiment 2A: One alcohol re-exposure session in a different context did not affect reinstatement

Following Pavlovian conditioning and extinction in Context A, rats received one alcohol re-exposure session in either Context A (‘Same’ group) or Context B (‘Different’ group), followed by a reinstatement test in Context A.

Counter to our predictions, conducting one alcohol re-exposure session in a different context had no effect on reinstatement. Relative to extinction, ΔCS port entries (Figure 4A) significantly increased at test [Phase: *F*_(1, 20)_ = 34.65, *p* < .001] in both groups [Group: *F*_(1, 20)_ = 0.04, *p* = .837; Phase x Group: *F*_(1, 20)_ = 0.01, *p* = .941]. The total duration of CS port entries (Figure 4B) significantly increased at test [Phase: *F*_(1, 20)_ = 17.01, *p* < .001] in both groups [Group: *F*_(1, 20)_ = 0.22, *p* = .647; Phase x Group: *F*_(1, 20)_ = 2.19, *p* = .155]. The average latency to initiate the first CS port entry (Figure 4C) significantly decreased at test [Phase: *F*_(1, 20)_ = 32.83, *p* < .001] in both groups [Group: *F*_(1, 20)_ = 0.83, *p* = .374; Phase x Group: *F*_(1, 20)_ = 0.13, *p* = .723].

**Figure 4.**
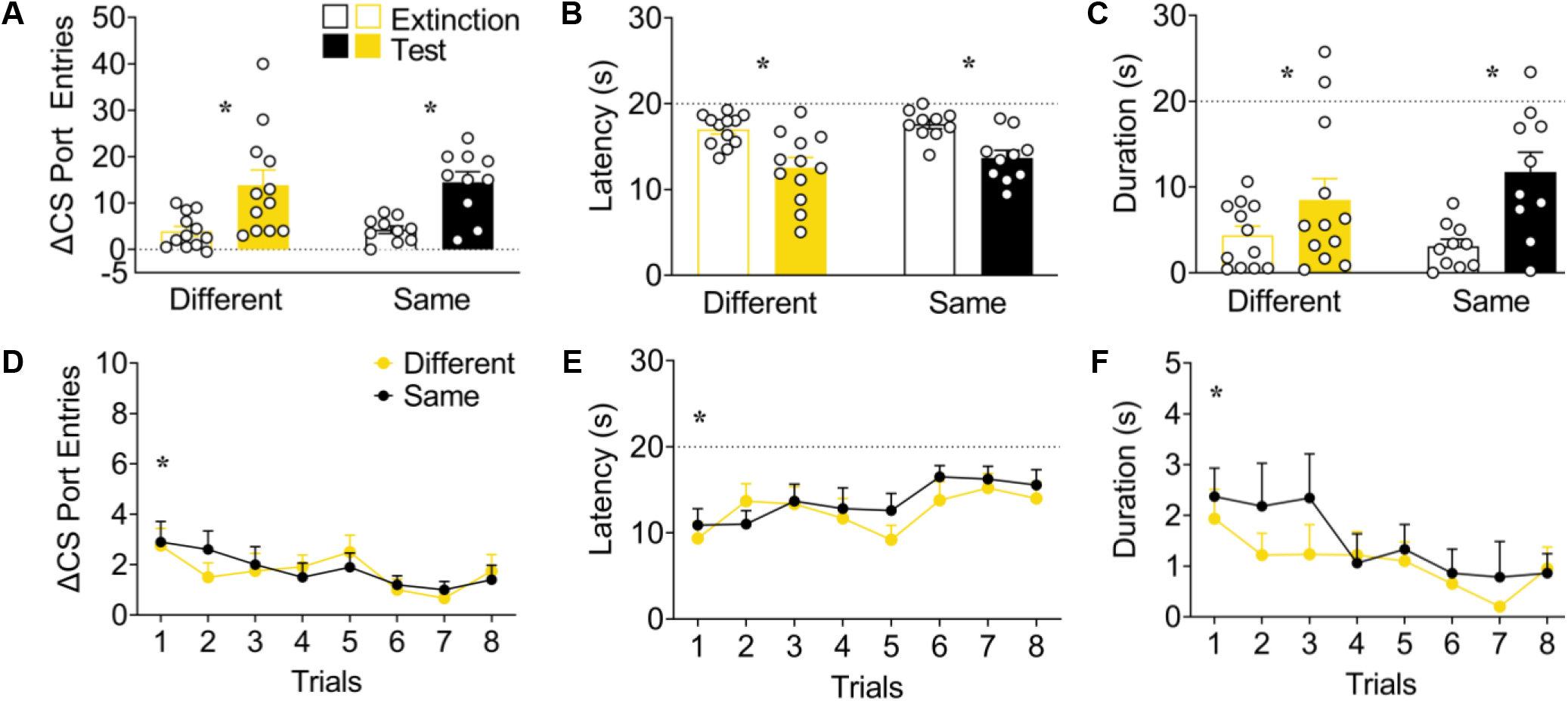
Conducting one alcohol re-exposure session in a different context did not reduce reinstatement. Data are from rats that received one alcohol re-exposure session in Context B (Different; Yellow; *n* = 12) or Context A (Same; Black; *n* = 10). All rats were tested in Context A. **A – C** Mean (± SEM) responding during extinction and test for **(a)** ΔCS port entries, **(b)** total duration of CS port entries and **(c)** average latency to initiate the first CS port entry. **D – F** Mean (± SEM) responding across CS trials at test for **(d)** ΔCS port entries, **(e)** duration of CS port entries, and **(f)** latency to initiate the first CS port entry. * *p* < 0.05, main effects of **A – C** Phase (Extinction vs. Test) and **D – F** Trial (1 – 8).

At test, ΔCS port entries (Figure 4D) significantly decreased across CS trials due to within session extinction [Trial *F*_(7, 140)_ = 3.24, *p* = .003] in both groups [Group: *F*_(1, 20)_ = 0.03, *p* = .874; Trial x Group: *F*_(7, 140)_ = 0.60, *p* = .754]. The total duration of CS port entries (Figure 4E) also significantly decreased across CS trials [Trial: *F*_(5.249, 104.984)_ = 3.07, *p* = .011] in both groups [Group: *F*_(1, 20)_ = 0.91, *p* = .351; Trial x Group: *F*_(5.249, 104.984)_ = 0.50, *p* = .786]. The average latency to initiate the first CS port entry (Figure 4F) significantly increased across CS trials [Trial: *F*_(7, 140)_ = 2.83, *p* = .009] in both groups [Group: *F*_(1, 20)_ = 0.54, *p* = .470; Trial x Group: *F*_(7, 140)_ = 0.56, *p* = .786]. These results show that conducting one alcohol re-exposure session in a different context did not affect reinstatement.

### Experiment 2B: Three alcohol re-exposure sessions in a different context reduced reinstatement during the first CS trial

Rats from Experiment 2A received additional Pavlovian conditioning and extinction in Context A, followed by three alcohol re-exposure sessions in either Context A (‘Same’ group) or Context B (‘Different’ group), then a reinstatement test in Context A.

Conducting three alcohol re-exposure sessions in a different context reduced reinstatement during the first CS trial. An analysis conducted on data averaged across the full test session showed that relative to extinction, ΔCS port entries (Figure 5A) significantly increased at test [Phase: *F*_(1, 21)_ = 56.29, *p* < .001] in both groups [Group: *F*_(1, 21)_ = 0.02, *p* = .882; Phase x Group: *F*_(1, 21)_ = 0.29, *p* = .597]. The total duration of CS port entries (Figure 5B) also significantly increased at test [Phase: *F*_(1, 21)_ = 27.01, *p* < .001] in both groups [Group: *F*_(1, 21)_ = 0.54, *p* = .472; Phase x Group: *F*_(1, 21)_ =1.46, *p* = .240]. The average latency to initiate the first CS port entry (Figure 5C) significantly decreased at test [Phase: *F*_(1, 21)_ = 91.16, *p* < .001] in both groups [Group: *F*_(1, 21)_ = 0.01, *p* = .929; Phase x Group: *F*_(1, 21)_ = 1.26, *p* = .274].

**Figure 5.**
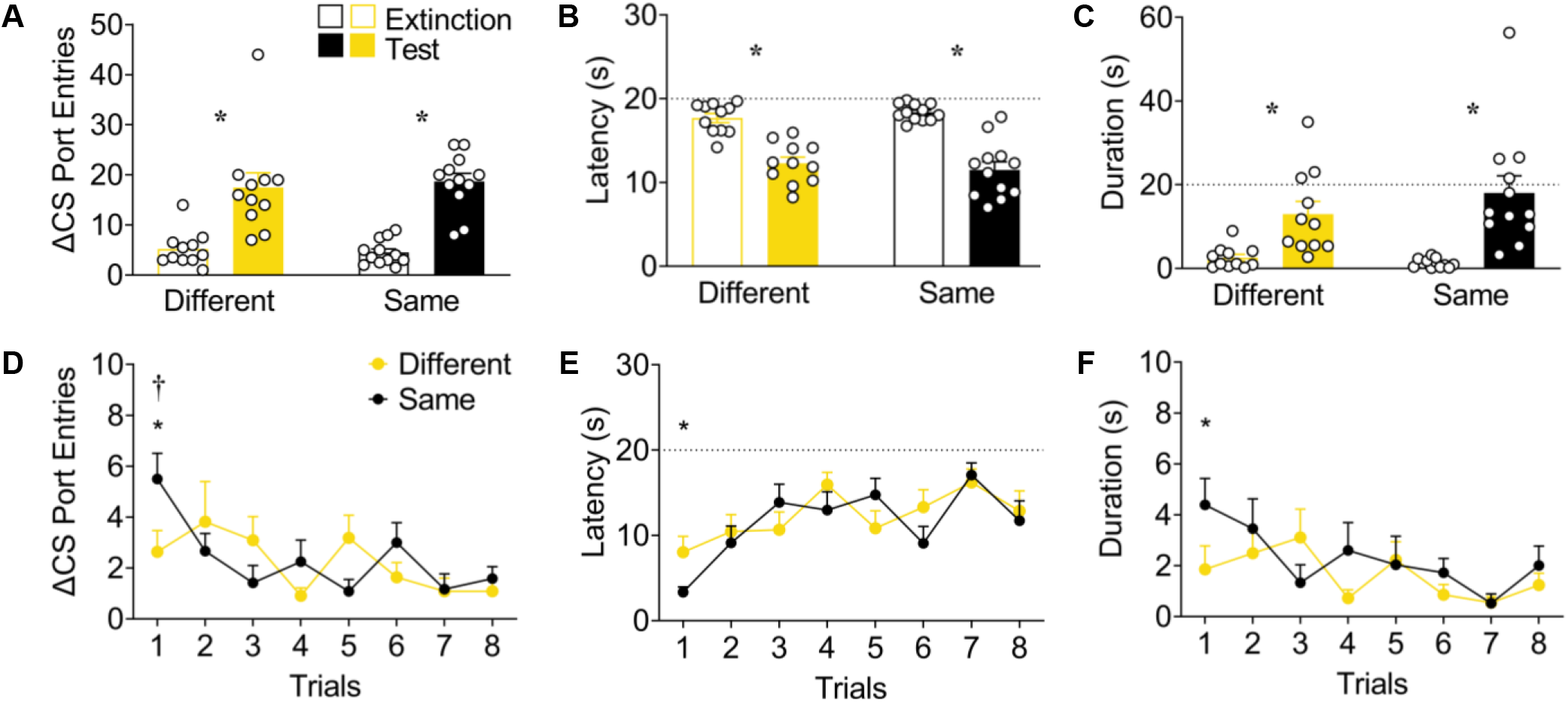
Conducting three alcohol re-exposure sessions in a different context reduced reinstatement during the first CS trial at test. Data are from rats that received three alcohol re-exposure sessions in Context B (Different; Yellow; *n* = 11) or Context A (Same; Black; *n* = 12). All rats were tested in Context A. **A - C** Mean (± SEM) responding during extinction and test for **(a)** ΔCS port entries, **(b)** total duration of CS port entries and **(c)** average latency to initiate the first CS port entry. **D - F** Mean (± SEM) responding across CS trials at test for **(d)** ΔCS port entries, **(e)** duration of CS port entries and **(f)** latency to initiate the first CS port entry. * *p* < 0.05, main effects of **A – C** Phase (Extinction vs. Test) and **D – F** Trial (1 – 8). † *p* < 0.05, Group x Trial interaction post-hoc (Different vs. Same on CS trial 1)

At test, ΔCS port entries (Figure 5D) significantly decreased across CS trials [Trial: *F*_(7, 147)_ = 3.26, *p* = .003]. Although there was no effect of group [Group: *F*_(1, 21)_ = 0.14, *p* = .717], ΔCS port entries differed between groups as a function of CS trial [Trial x Group: *F*_(7, 147)_ = 2.44, *p* = .022]. Post-hoc analyses revealed that the ‘Different’ group made significantly fewer ΔCS port entries during the first CS trial, compared to the ‘Same’ group (*p* = .042). The total duration of CS port entries (Figure 5E) and average latency to initiate the first CS port entry (Figure 5F) did not, however, follow this pattern of responding. The total duration of CS port entries significantly decreased across CS trials [Trial: *F*_(6.269, 131.655)_ = 2.81, *p* = .012] in both groups [Group: *F*_(1, 21)_ = 0.95, *p* = .340; Trial x Group: *F*_(6.269, 131.655)_ = 1.67, *p* = .130]. The average latency to initiate the first CS port entry (Figure 5F) significantly increased across CS trials [Trial: *F*_(7, 147)_ = 6.31, *p* < .001] in both groups [Group: *F*_(1, 21)_ = 0.42, *p* = .524; Trial x Group: *F*_(7, 147)_ = 1.57, *p* = .150]. Thus, conducting three alcohol re-exposure sessions in a different context modestly reduced reinstatement, as seen by a reduction in ΔCS port entries during the first CS trial.

## Discussion

This study examined the psychological processes involved in the reinstatement of responding to an appetitive, alcohol-predictive CS. We found that spaced context extinction did not affect reinstatement, whereas massed context extinction prevented reinstatement. Moreover, conducting one alcohol re-exposure session in a context that differed from the subsequent test context had no effect on reinstatement, whereas conducting three alcohol re-exposure sessions in a different context from the test context reduced the reinstatement of port entries during the first CS trial. These findings partially support a view generated from aversive Pavlovian conditioning procedures, which posits that a context-US association formed during US re-exposure plays a role in reinstatement.

In Experiment 1A, we extinguished the context-alcohol association formed during the alcohol re-exposure session by exposing rats to the conditioning chambers that alcohol re-exposure was conducted in across four daily sessions (‘ spaced context extinction’). Total port entries decreased across sessions in this group, suggesting that context extinction had occurred. Control groups were exposed either to the conditioning chambers without the fluid ports or to an alternate context for the same duration.

Counter to our expectations, all three groups showed significant reinstatement as measured by ΔCS port entries, CS port entry duration, and CS port entry latency. These results differ from evidence that a comparable, spaced context extinction manipulation diminished the reinstatement of operant food-seeking (Baker et al., 1991). Thus, a context-US association formed during US re-exposure may be involved in the reinstatement of operant responding for a food reinforcer, but not in the reinstatement of Pavlovian responding to an alcohol-predictive CS. Alternatively, our context extinction procedure may not have sufficiently extinguished the context-alcohol association, despite producing a decrease in total port entries across context extinction sessions.

To evaluate the latter possibility, we tested the effect of massed context extinction on reinstatement. Given that presenting CS trials in a temporally massed manner extinguishes conditioned responding to an aversive CS to a greater degree than temporally spaced CS trials (Cain et al., 2003), we hypothesized that conducting massed extinction sessions would deepen context extinction and reduce reinstatement. In Experiment 1B, rats were exposed to either the context that alcohol re-exposure occurred in or to an alternate context in four sessions delivered across two days. A third group that did not receive alcohol re-exposure and was then exposed to either the training or alternate context served as a control for the potential spontaneous recovery of CS port entries after extinction. Interestingly, massed context extinction after alcohol re-exposure significantly reduced reinstatement, as indexed by ΔCS port entries and latency to make the first CS port entry, but not duration of CS port entries. Moreover, rats that did not receive alcohol re-exposure did not show changes in conditioned responding at test relative to extinction, indicating that spontaneous recovery did not contribute to reinstatement in our task. These results concur with previous work showing that a similar massed context extinction manipulation reduced reinstatement to an aversive CS (Bouton & Bolles, 1979), and support the view that a context-US association formed during US re-exposure plays a role in reinstatement to an appetitive, alcohol-CS (Bouton & Bolles, 1979).

In Experiment 2A, we determined if conducting alcohol re-exposure in Context B, that differed from the subsequent test context (Context A), would impact reinstatement. Counter to our predictions, there was no effect of this manipulation on reinstatement as measured by ΔCS port entries, CS port entry duration, and CS port entry latency. These findings contradict studies showing that reinstatement to an aversive CS did not occur when US re-exposure was conducted in a context that differed from the test context (Bouton, 1984; Bouton & Bolles, 1979; Bouton & King, 1983; Frohardt et al., 2000; Wilson et al., 1995). To address the possibility that one alcohol re-exposure session may not have been sufficient for the rats to associate Context B with alcohol, in Experiment 2B we tested the effect of conducting three alcohol re-exposure sessions in Context B on subsequent reinstatement in Context A. This manipulation had no effect on reinstatement as assessed by data averaged across the full test session. However, it significantly reduced ΔCS port entries during the first CS trial at test. Although a seemingly modest effect, in aversive Pavlovian conditioning tasks learning is sometimes assessed after just one CS-US trial (Han et al., 2008; Park et al., 2020; Vieira et al., 2014), and responding at test is sometimes assessed in one or two CS trials (Bouton & Bolles, 1979; Han et al., 2008; Lebrón, Milad, & Quirk, 2004). Arguably, conditioned responding elicited by the first CS trial at test may be the best indicator of an animal’s expectation regarding whether or not the US will occur.

Several published studies conducted using aversive conditioning procedures have shown that conducting US re-exposure in a context that differs from the test context prevents reinstatement (Bouton, 1984; Bouton & Bolles, 1979; Bouton & King, 1983; Frohardt et al., 2000; Wilson et al., 1995). One interpretation of these data is that reinstatement does not occur because the context-US association formed during US re-exposure is not present at test to summate with the residual predictive value of the CS that survived extinction to drive reinstatement. An alternate hypothesis regarding the reinstatement effect is that reinstatement may be due to the US re-exposure session reactivating the CS representation, as the context can become associated with the CS during previous extinction training. This CS representation would be experienced in tandem with the US delivery during the US re-exposure session, which could result in a strengthened CS-US association and drive reinstatement 24 h later (Holland, 1990; Westbrook et al., 2002). If the US re-exposure session were conducted in a different context, which was never associated with the CS through extinction training, the CS representation would not be activated. Therefore, the CS representation would not become associated with the US and would not produce subsequent reinstatement. According to either hypothesis, we should not have observed reinstatement when alcohol re-exposure was conducted in an alternate context. Interestingly, prior data has shown that re-exposure to a food-US in either the test context or a different context reinstated conditioned responding to a food-predictive CS; however, reinstatement was more robust when food re-exposure occurred in the same context as the subsequent reinstatement test (Bouton & Peck, 1989). Therefore, it is not surprising that we found some reinstatement of CS port entries at test following three alcohol re-exposure sessions in Context B.

In Experiment 2, reinstatement following alcohol re-exposure in Context B may have been a result of context generalization. The alternate (Context B) and training (Context A) contexts differed in terms of sensory stimuli (i.e., odour, tactile, visual); however, the innate features of the conditioning chambers were consistent across contexts (i.e., house light, speaker, and fluid port position). These similarities could have resulted in a context-alcohol association formed in Context B generalizing to the test context. Future studies could address this possibility by conducting context preference tests after alcohol re-exposure. If rats that are re-exposed to alcohol in Context B show a similar preference for the alternate and test contexts, this could suggest that the context-US association has generalized across the two distinct contexts. Another consideration with the experimental design of Experiment 2 is that the same groups were used in Experiments 2A and 2B. Therefore, the reduction in reinstatement seen during CS trial 1 in the ‘ Different’ group may have been the result of repeated testing under the same conditions. Alternatively, it is possible that more sessions of alcohol re-exposure were needed to fully unmask an effect on reinstatement.

A unique aspect of appetitive conditioning tasks is that a context-US, or strengthened CS-US, association may not be the sole mechanisms contributing to reinstatement. A consummatory response is required to voluntarily ingest an appetitive US like alcohol and this consummatory response may contribute to reinstatement. This possibility is supported by our previous work showing that re-exposure to water as a control condition reinstated responding to an alcohol-predictive CS to the same degree as re-exposure to alcohol, whereas when alcohol re-exposure occurred via systemic injection reinstatement was not observed (LeCocq et al., 2018). This additional factor of a consummatory response does not occur in aversive conditioning tasks; therefore, this difference could account for discrepancies in our findings and previous findings using aversive conditioning procedures. Future studies could assess the impact of consummatory behaviour on the reinstatement of responding to an alcohol-predictive CS by delivering alcohol during the alcohol re-exposure session in a manner that produces a different consummatory response from that used during Pavlovian conditioning (e.g., via a sipper tube instead of in the fluid port).

Finally, an important consideration in the present research is that we only used male rats. Given the generality of the reinstatement effect and its importance in evaluating the role of cues in people with substance use disorders or post-traumatic stress disorders, it is critical for preclinical research to be conducted using both male and female subjects (Radke, Sneddon, & Monroe, 2021; Sanchis-Segura & Becker, 2016; Shansky, 2015; Shansky & Murphy, 2021). Our recent, unpublished data show that female rats reinstate responding to a sucrose-predictive CS to a greater degree relative to male rats (LeCocq & Chaudhri, 2021), and we are currently using male and female rats in ongoing experiments to study the role of μ-opioid receptors in reinstatement (LeCocq & Chaudhri, 2021).

In conclusion, our findings extend the literature on the psychological processes underlying reinstatement by providing evidence that a context-US association plays a role in the reinstatement of responding to an alcohol-predictive CS. However, more research is needed to elucidate the role that consummatory behaviours may play in this reinstatement effect. Thus, these findings provide the basis for future studies aimed at investigating processes that may uniquely underlie appetitive reinstatement.

## Acknowledgements

This research was funded by the Canadian Institution of Health Research (MOP-137030, N.C.). N. C. was the recipient of a Chercheur-Boursier award from the Fonds de la recherche en santé Québec and is a member of the Center for Studies in Behavioral Neurobiology. M. R. L. was supported by a graduate scholarship from the Faculty of Arts and Science at Concordia University and by a Fonds de recherche du Québec - Nature et technologies doctoral fellowship. S.S. was supported by a Concordia Undergraduate Summer Research Award. The authors would like to thank Stephen Cabilio for support with Med-PC programming and data extraction. All authors confirm that there are no conflicts of interest.

