## Supplementary material for "The Role of Context Conditioning in the Reinstatement of Responding to an Alcohol-Predictive Conditioned Stimulus"

**Supplementary Table 1.** Number of rats receiving different alcohol concentrations during Pavlovian conditioning and alcohol re-exposure sessions

| **Experiment** | **Number of rats on 5% EtOH** | **Number of rats on 15% EtOH** |
| --- | --- | --- |
| 1A | *n* = 0 | *n* = 36 |
| 1B | *n* = 0 | *n* = 36 |
| 2A | *n* = 1 | *n* = 27 |
| 2B | *n* = 1 | *n* = 27 |

| **Intermittent alcohol access** | | | |
| --- | --- | --- | --- |
| **A** | 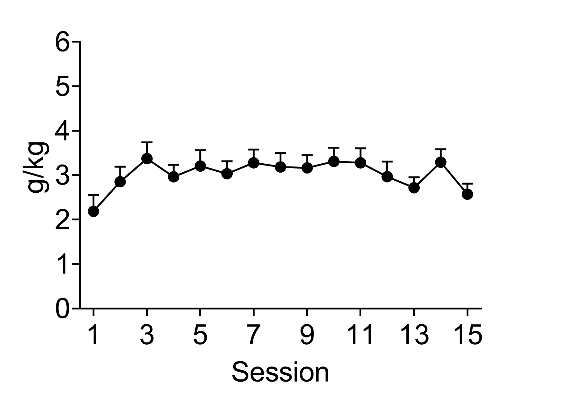 | **B** | 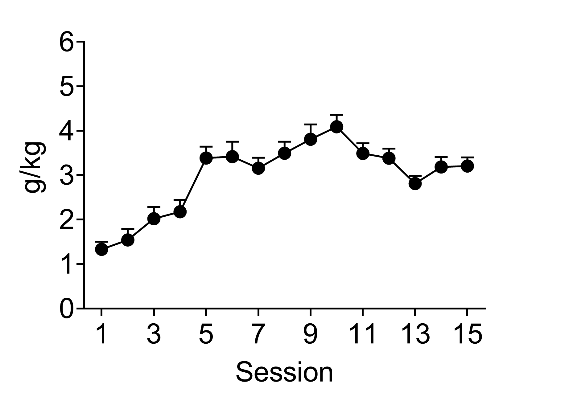 |

**Supplementary Figure 1.** A repeated measures ANOVA revealed that alcohol intake (g/kg) increased across intermittent alcohol access sessions in **A** Experiment 1 [*F*_(6.725, 235.358)_ = 16.343, *p* < 0.001], and **B** Experiment 2 [*F*_(5.291, 142.856)_ = 2.276, *p* = 0.047]. Data represent mean (± SEM) g/kg obtained across sessions.

| **Acquisition and extinction of conditioned responding for alcohol** | | | |
| --- | --- | --- | --- |
|  | **Experiment 1A** |  | **Experiment 2A** |
| **A** | 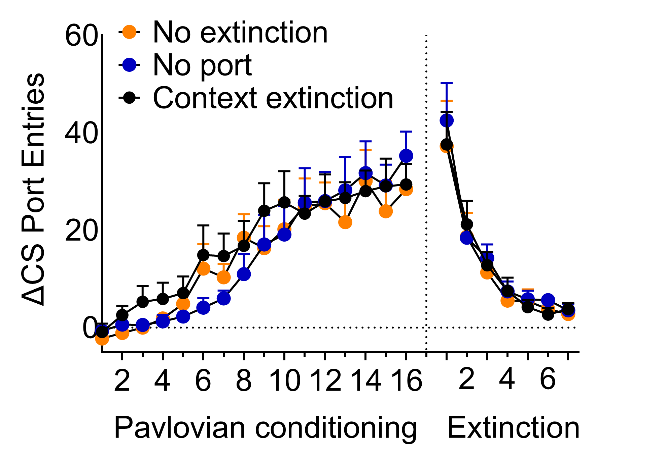 | **B** | 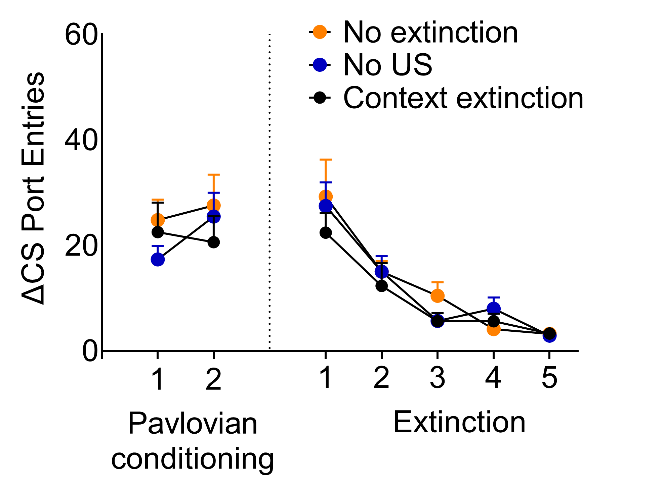 |
|  | **Experiment 2A** |  | **Experiment 2B** |
| **C** | 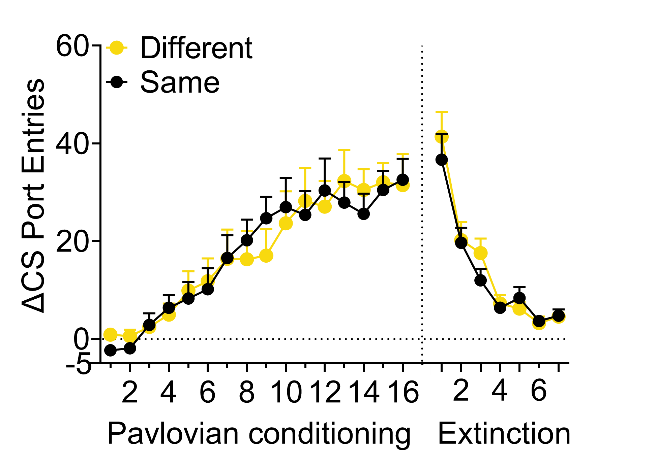 | **D** | 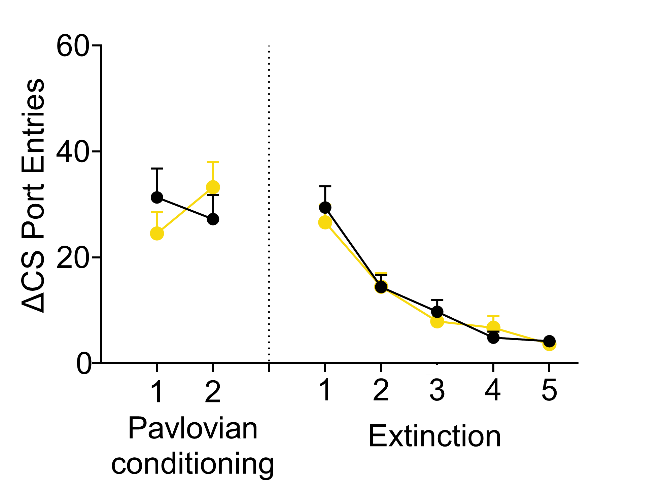 |

**Supplementary Figure 2.** Rats learned to associate the CS with alcohol in both experiments. Separate Session x Group mixed ANOVA revealed that ΔCS port entries significantly increased across Pavlovian conditioning sessions similarly in all groups in **A** Experiment 1A [Session: *F*_(4.380, 118.266)_ = 38.760, *p* < .001; Session x Group: *F*_(8.760, 118.266)_ = 0.847, *p* = 0.572; Group: *F*_(2, 27)_ = 0.206, *p* = 0.815], **B** Experiment 1B [Session: *F*_(1, 25)_ = 2.076, *p* = 0.162; Session x Group: *F*_(2, 25)_ = 1.966, *p* = 0.161; Group *F*_(2, 25)_ = 0.374, *p* = 0.692], **C** Experiment 2A [Session: *F*_(4.472, 89.449)_ = 25.057, *p* <0.001; Session x Group: *F*_(4.472, 89.449)_ = 0.523, *p* = 0.739; Group: *F*_(1, 20)_ = 0.000, *p* = 0.984], and **D** Experiment 2B [Session: *F*_(1, 21)_ = 0.431, *p* = 0.518; Session x Group: *F*_(1, 21)_ = 3.283, *p* = 0.084; Group: *F*_(1, 21)_ = 0.004, p = 0.947].

The conditioned response extinguished, as a separate Session x Group mixed ANOVA revealed that ΔCS port entries significantly decreased across extinction sessions similarly in all groups in **A** Experiment 1A [Session: *F*_(2.143, 57.869)_ = 48.234, *p* < .001; Session x Group: *F*_(4.287, 57.869)_ = 0.195, *p* = .948; Group: *F*_(2, 27)_ = 0.166, *p* = 0.848], **B** Experiment 1B [Session: *F*_(2.447, 61.165)_ = 40.046, *p* < 0.001; Session x Group: *F*_(4.893, 61.165)_ = 0.676, *p* = 0.640; Group: *F*_(2, 25)_ = 0.490, *p* = 0.618], **C** Experiment 2A [Session: *F*_(2.855, 57.108)_ = 57.713, *p* < 0.001; Session x Group: *F*_(2.855, 57.108)_ = 0.724, p = 0.535; Group: *F*_(1, 20)_ = 0.259, *p* = 0.616], and **D** Experiment 2B [Session: *F*_(2.864, 60.151)_ = 41.413, *p* < 0.001; Session x Group: *F*_(2.864, 60.151)_ = 0.355, *p* = 0.777; Group: *F*_(1, 21)_ = 0.121, *p* = 0.732]. Data represent mean (± SEM) ΔCS port entries across sessions.
